# Size-dependent movement explains why bigger is better in fragmented landscapes

**DOI:** 10.1101/264861

**Authors:** Jasmijn Hillaert, Thomas Hovestadt, Martijn L. Vandegehuchte, Dries Bonte

**Author notes:** **Corresponding author** Jasmijn Hillaert Terrestrial Ecology Unit Department of Biology Ghent University K. L. Ledeganckstraat 35 9000 Ghent Belgium Corresponding author.

## Abstract

Body size is a fundamental trait known to allometrically scale with metabolic rate, and therefore a key determinant of individual development, life history and consequently fitness. In spatially structured environments, movement is an equally important driver of fitness. Because movement is tightly coupled with body size, we expect habitat fragmentation to induce a strong selection pressure on size variation across and within species. Changes in body size distributions are then, in turn, expected to alter food web dynamics. However, no consensus has been reached on how spatial isolation and resource growth affect body size distributions.

Our aim was to investigate how these two factors shape the body size distribution of consumers under scenarios of size-dependent and -independent consumer movement by applying a mechanistic, individual-based resource-consumer model. The outcome was then linked to important ecosystem traits such as resource abundance and stability. Finally, we determined those factors that explain most variation in size distributions.

We demonstrate that decreasing connectivity and resource growth select for communities (or populations) consisting of larger species (or individuals) due to strong selection for the ability to move over longer distances. When including size-dependent movement, moderate levels of connectivity result in increases in local size diversity. Due to this elevated functional diversity, resource uptake is optimized at the metapopulation or metacommunity level. At these intermediate levels of connectivity, size-dependent movement explains most of the observed variation in size distributions. Interestingly, local and spatial stability of consumer biomass are lowest when isolation and resource productivity are high. Finally, we highlight that size-dependent movement is of vital importance for the survival of populations within highly fragmented landscapes. Our results demonstrate that considering size-dependent movement and resource growth is essential to understand patterns of size distributions at the population or community level and the resulting metapopulation or metacommunity dynamics.

## Introduction

Body size sets limits to the functioning of individuals, thereby affecting inter- and intraspecific interactions and regulating overall food web structure (Bartholomew et al. 1982, Brose et al. 2006). Body size is also central to metabolic theory (Kleiber 1932, Brody et al. 1934, Brody 1945, Brown et al. 2004). Starting from the simple allometric rule linking body size with metabolic rate, important inferences can be made at the level of individuals, populations, communities, and ecosystems (Brown et al. 2004). For instance, the ingestion rate and speed of movement of an individual are correlated with its body size (Peters 1983). Also, shifts in community size structure have been shown to affect ecosystem functioning (Yvon-Durocher and Allen 2012, Fritschie and Olden 2016). Hence, body size can be considered a super trait (Fritschie and Olden 2016, Brose et al. 2017) relating to both ecological effects and responses and therefore constraining ecological and evolutionary dynamics (Applebaum et al. 2014, Llandres et al. 2015).

Body size is directly related to individual biomass. With increasing resource productivity, more total consumer metabolic biomass can be supported (Atkins et al. 2015). However, for a given amount of resources, higher abundance implies lower per capita energy use (i.e., the energetic equivalence rule). Therefore, increased productivity can either result in more or larger individuals (Damuth 1981, White et al. 2007, Ehnes et al. 2014). Furthermore, resources are usually not homogeneously distributed across space, but spatially structured (Krummel et al. 1987). This implies that organisms need to move both within (foraging resulting in spatially coupled patches) and across generations (dispersal resulting in metapopulation dynamics) to make optimal use of resources, depending on the spatiotemporal dynamics of both the resource and the consumer population (Amarasekare 2008). Because body size determines to a large extent the movement capacity of active dispersers (Stevens et al. 2014), we expect it to have a large impact on population and community dynamics in spatially structured environments and to be under strong selection.

The cost of movement is highly dependent on resource availability and habitat connectivity, and is one of the costs that changes with body size (Peters 1983, Bonte et al. 2012). Large-sized individuals may, for instance, incur higher costs due to their larger home ranges but more directly, we expect larger body sizes to be associated with reduced time costs because of higher achieved speed and increased perceptual range (Buddenbrock 1934, Peters 1983, Mech and Zollner 2002, Pawar et al. 2012). This view is at the basis of the textural discontinuity hypothesis, which states that the modes of a size abundance distribution mirror those scales at which resources within the landscape are most abundant, relative to the size and dispersal capacity of the consumer species (Holling 1992, Borthagaray et al. 2012, Nash et al. 2014). Despite the attention of theoretical studies to the origin of size distributions (e.g. Loeuille & Loreau 2005; Ritterskamp, Bearup & Blasius 2016), only few have covered the dependence of size distributions on habitat configuration (but see Milne *et al.* 1992; Etienne & Olff 2004; Borthagaray *et al.* 2012; Buchmann *et al.* 2012, 2013). Overall, theory based only on spatial scaling of size versus resource distribution predicts that resource availability and distribution strongly affect body size distributions of species within communities (Holling 1992, Ritchie and Olff 1999, Allen et al. 2006, Borthagaray et al. 2012, Nash et al. 2014).

Body size is not only central to mobility and metabolic rate, but also to development (West et al. 2001). Small individuals and species have the advantage of low energy requirements and short developmental times (Peters 1983). Large individuals and species, on the other hand, are capable of crossing unsuitable matrix to reach new patches and have higher tolerances to starvation (Peters 1983, Davies et al. 2000, Tscharntke and Brandl 2004). This could explain why, although many empirical studies have investigated the effect of habitat fragmentation on body size distributions, a conclusive pattern remains elusive (e.g. (Thomas 2000a, Davies et al. 2000, Hamback et al. 2007, Jauker et al. 2016, Warzecha et al. 2016, Renauld et al. 2016). It thus remains difficult to predict how the spatial distribution of resources affects body size distributions.

Due to the ubiquitous increase in habitat loss and fragmentation we urgently need to better understand communities’ and species’ responses to isolation. Fragmentation studies on mammals and birds or migratory behavior in insects are executed at large scales (Thomas 2000; Gonzalez & Chaneton 2002; Debinski & Holt 2009; Haddad *et al.* 2015). However, little research is performed on the effect of fragmentation and isolation at the small scales at which insects forage (Gonzalez and Chaneton 2002, Braschler and Baur 2016).

As body size is anticipated to be central to both movement and resource consumption, its distribution in space and time will have a strong impact on ecosystem stability, primary productivity, and biodiversity (Massol et al. 2017). Individuals in metapopulations or metacommunities function as mobile linkers that organize themselves in space to maximize their fitness according to their size (Jeltsch et al. 2013). Further, stabilizing mechanisms allow for species coexistence, increasing diversity, which has been shown to be positively affected by habitat fragmentation (Jeltsch et al. 2013, Arnillas et al. 2017, Fahrig 2017). Because of the prominent feedback of consumer variability and mobility on lower trophic levels, it is expected that other ecosystem traits such as resource abundance are strongly affected by level of connectivity and size variation. Resource abundance will then again alter consumer biomass, regulating ecosystem stability which is crucial for ecosystem functioning and sustainability and varies across scales (Wang and Loreau 2014). Because current theory fails to formally link selection on body size to metabolic and metapopulation theory, we created an individual-based, spatially explicit model to study the effect of fine-grained resource isolation on the selection of body size distributions of a consumer species or community. As habitat isolation might affect a resource’s growth speed by changing abiotic factors, we also test how resource growth speed interacts with habitat isolation in affecting the selection of body size. Based on established allometric rules linking body size to movement speed, movement costs, basal metabolic rate, ingestion rate, growth rate, and reproduction we created a mechanistic model, to increase our understanding of how differences in the isolation and productivity of resources affect a consumer’s body size distribution. The development of such a complex, mechanistic model is a necessity when studying body size as these allometric rules imply the existence of crucial trade-offs, which should not be overlooked. We further aim to uncover the importance of size-dependent movement for selection and ecological dynamics. Moreover, the impact on crucial ecosystem traits such as ecosystem stability at various scales and resource abundance is estimated. Our individual-based approach enables us to interpret our results either as within-species adaptive dynamics of individual body size or across-species metacommunity changes in the distribution of species of different size. Overall, we expect an increasing importance of size-dependent movement in environments with increasing isolation of the resources and thus a (community-wide) shift towards larger body sizes as habitat fragmentation increases.

## Methodology

We developed a consumer-resource model to understand how the spatial distribution of a resource affects the size distribution of its consumer(s). The spatial distribution of the resource and its abundance differed between simulations with regard to the distance between its suitable patches (*NND*) and its growth rate. This resource may be consumed by (a community of) consumers. All traits of the consumers are related to their mass by allometric rules, as derived from literature (e.g. Peters 1983). An individual’s body mass is used to represent its size (Peters 1983). Also, we assessed the importance of size-dependent movement for shaping the evolved consumer size distribution and its impact on metapopulation functioning. This was done by creating two models: a coupled and a decoupled model. In the coupled model, speed of movement and perceptual range both increase with body size, whereas in the decoupled model, body size, perceptual range and speed of movement are unlinked.

The model is a spatially explicit, discrete-time model with overlapping generations. One time step corresponds to one day within the lifetime of the consumer. We here took an arthropod-centered angle and parameterized allometric rules for a haploid, parthenogenetic arthropod species feeding on plants (the resource) with a semelparous lifecycle. All parameters of the model are summarized in Table S4.1.

### The landscape

The landscape is cell-based with each cell having a side length (*SL*) of 0.25 m. This fine-grained fragmentation is relevant for studying the effect of isolation on the foraging behavior of arthropods. Within the landscape, a distinction is made between suitable and unsuitable habitat. Resources only grow within suitable patches with one patch having the size of a single cell. All landscapes have a constant number of suitable patches (i.e. 2500) but varying nearest neighbor distance (*NND*) (Fahrig 2003). The effect of isolation is tested by assigning a constant *NND* from 0 to 10 to all cells (see supplementary material part 3 for an example). Consequently, the dimensions of the landscape increase with *NND* according to (50 + *NND*50*) × (50 + *NND*50*) cells. The boundaries of the landscape are wrapped.

### The resource

As it is advisable not to focus on individual species but also cover their interactions with other species (Berg et al. 2010), we included the dependence of the consumer on its resource by varying the resource’s growth speed. Resources at the cell level are not individually modeled but by a local logistic growth model. Local resource biomass is represented as the total energetic content of resource tissue within that cell (*R_x,y_* in joules). This resource grows logistically in time depending on the resource’s carrying capacity (*K*) and intrinsic growth rate (*r). K* was set to 2000 *J* [assumption of space limitation] whereas *r* differed between simulations (0.1, 0.5 or 0.9 per day; assumption for productivity of the system). In any cell, a fixed amount of resource tissue *[E_nc_*, in Joules, fixed at *1 J*) is non-consumable by the consumer species, representing below-ground plant parts. As such, *E_nc_* is the minimum amount of resource tissue present within a suitable cell, even following local depletion by the consumer species.

### The consumer

All consumers are individually modeled within the landscape. The consumer species has two life stages: a juvenile and adult life stage. Within each time step (a day), both stages have the chance to execute several events (see Figure 1).

**Figure 1:**
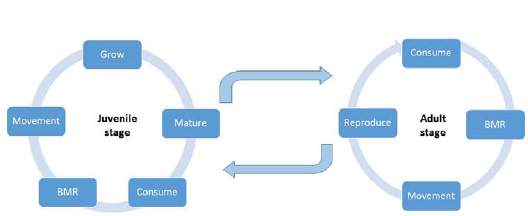
A comparison of daily events for the juvenile and adult stage of the consumer. BMR stands for the basal metabolic rate costs.

First, all individuals nourish their energy reserve by consumption. An individual’s energy reserve is depleted again by the cost of daily maintenance (i.e. basal metabolic rate) and movement. Only juveniles have the chance to grow and mature, whereas adults are able to reproduce, both of which further deplete the energy reserve. As the consumer species is semelparous, adults die after reproduction. How body size is implemented within each of these events is explained in supplementary material part 4.

Individual body size at maturity (*W_max_*, in kg) is coded by a single gene. Adult size is heritable and may mutate with a probability of 0.001 during reproduction. A new mutation is drawn from the uniform distribution [*W_max_* - [*W_ma_*/2), *W_max_* + [*W_max_*/2)] with *W_max_* referring to the adult size of the parent. New mutations must not exceed the predefined boundaries [0.01g, 3g] that represent absolute physiological limits. As such, our minimum adult size corresponds to the size of a small grasshopper such as *Tetrix undulata* (0.01 g) and the maximum size (3 g) to that of some longhorn beetles (Cerambycidae), darkling beetles (Tenebrionidae), scarab beetles (Scarabaeidae) or grasshoppers (Acrididae). New variants of this trait might also originate from immigration. The former source of variation enables fine-tuning of the optimal body size, whereas the latter source of variation facilitates fitness peak shifts.

### The movement phase

#### Emigration rate

Whether an individual moves depends on the ratio of the amount of energy present within a cell (R_x,y_) relative to the maximum amount of energy that can be consumed by all consumers present within that cell. This latter factor is determined by calculating the sum of all individuals’ daily ingestion rates within that cell 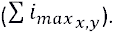

By assuming a symmetric competition, the probability of moving (*p*) is equal for all individuals present within the same cell and is calculated by (based on (Poethke and Hovestadt 2002)):

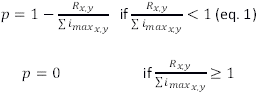

#### Determining cell of destination

As one time step in our model corresponds to one day, we do not model the movement behavior of an individual explicitly but instead, estimate the total area an individual can cover during a day in search for resources. This total area an individual can search during a day is called its foraging area, which is circular and is defined by a radius (*rad*, see further). The center of an individual’s foraging area corresponds to its current location. Overall, the size of an individual’s foraging area increases with its size (Peters 1983, Tscharntke and Brandl 2004) and is recalculated daily by taking into account an individual’s optimal speed (*v_opt_*), movement time (*t_m_*) and perceptual range (*d_per_*). During each movement event, an individual selects the patch with the highest amount of resources within its foraging area (i.e. informed settlement).

An individual’s optimal speed of movement (*v_opt_*, in meters per second) is calculated by means of the following equation, which has been based on walking insect (Buddenbrock 1934 in Peters 1983):

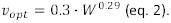

Here, *W* refers to the current weight of an individual, expressed in kg. The time an individual invests in movement per day (*t_m_*, in seconds) is maximally 1 hour. In case too little internally stored energy is present to support movement for one hour, *t_m_* is calculated by:

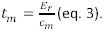

*E_r_* refers to the energy available within an individual’s energy reservoir and *c_m_* to the energetic cost of movement (in joules per second). The latter is calculated by the following formula which is based on running poikilotherms (Buddenbrock 1934 in Peters 1983):

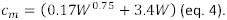

Independent of the cell of destination, the cost of moving during the time *t_m_ (t_m_ ·c*_m_) is subtracted from an individual’s energy reserve.

Based on *t_m_* and *v_opt_*, the total distance an individual covers at day *t* (*d_max_*) is determined as:

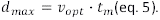

Next, the perceptual range of an individual is determined by means of the following relationship:

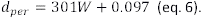

For simplicity, this relationship is linear and based on the assumption that the smallest individual (0.01g) has a perceptual range of 0.10 m and the largest individual (3g) a perceptual range of 1m. The effect of this relationship has been tested (see sensitivity analysis). Moreover, the positive relationship between body size and perceptual range has been illustrated over a wide range of taxa, including arthropods (supplementary information of (Pawar et al. 2012)).

Finally, the foraging area of an individual is circular and its radius (*rad*, in m) is calculated based on the total distance the individual has covered during the day and the individual’s perceptual range (see supplementary material part 2 for explanation of this formula):

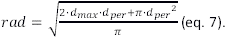

### Coupled versus decoupled model

To determine the importance of size-dependent movement, two different models were created: a coupled and a decoupled model. In the coupled model, speed of movement (*v_opt_*) and perceptual range both increase with body size. The decoupled model represents a null model in which body size, speed of movement and perceptual range were unlinked. Body size and speed of movement were unlinked by resampling an individual’s speed of movement each day from the uniform range [0.0106, 0.0557], Here, 0.0106 corresponds to the optimal speed of the smallest adult individual (0.01 g) and 0.0557 to the optimal speed of the largest adult individual (3 g). Also, the perceptual range of an individual is no longer increasing with body size, but instead sampled daily from the uniform distribution [0.1 m, 1 m]. We chose to sample from a uniform distribution rather than from an evolved scenario in the decoupled model to avoid any skewness and bias in the randomization. As the cost of movement is based on the total movement time and not total distance, it is unaffected by the decoupling.

### Data analysis

Within the coupled model, the simulations with *NND* 10 and growth speed 0.1 and 0.5 went extinct without immigration from outside the landscape, therefore, we omitted these simulations during the analysis.

During each simulation, we traced changes in the mean amount of resources per cell, total number of adults and larvae, average adult weight (*W_max_*) and the coefficient of variation, skewness, and kurtosis of the consumer’s adult weight (*W_max_*) distribution. Every 500 time steps, the value of *W_max_* of maximum 50 000 randomly sampled individuals was collected.

### Occupancy(O)

Occupancy (*O*) is defined as the ratio of occupied patches to the total number of suitable patches within the landscape. The level of occupancy is determined every ten days during the last 100 days of a simulation. At the end, the average of these values is calculated per simulation.

### Variability

In order to infer the stability of the community at several scales we calculated the *α, β_2_* and *γ* variability per simulation. α variability is a measure of the local temporal variability and is calculated by

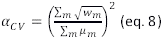

With *w_m_* referring to the temporal variance and *μ_m_* to the temporal mean of community consumer biomass in cell *m* (Wang and Loreau 2014). The temporal variability at the metacommunity scale or *y* variability was calculated by:

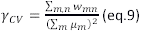

With *w_mn_* referring to the temporal covariance of community biomass between cells *m* and *n* (Wang and Loreau 2014). Finally, *β_2_* variability or asynchrony-related spatial variability was determined by:

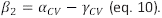

In order to calculate these variables, we recorded the total consumer biomass of 100 randomly selected suitable patches every ten days during the final 100 days of a simulation.

### Reproductive success and movement

Throughout the final 600 days of a simulation, 1000 eggs were randomly selected to be followed during their lifetime. Their movements and reproductive success were recorded.

### Variation partitioning

By means of multivariate variation partitioning we disentangled the amount of variation in adult size that can be explained by the coupling of body size and movement, resource growth rate and level of isolation. Analysis were performed in R by applying the function varpart within the package vegan which is based on calculating the adjusted *R^2^* in redundancy analysis ordination (RDA) (Oksanen et al. 2018). This was done by collecting the average, coefficient of variation, level of skewness, and level of kurtosis of the distribution of *W_max_* per simulation. We also executed a similar analysis for (i) occupancy, (ii) parameters summarizing resource and consumer dynamics (resource abundance, resource variance and consumer abundance) and (iii) the metapopulation functioning statistics *α*, *β*_2_ and *γ* variability. We executed a global variation partitioning including all distances except for *NND* 10 as some of these simulations were not stable and only survived as sinks. We furthermore executed a variation partitioning for each value of *NND* independently. As such, the effect of isolation on the amount of variation explained by the coupling of body size and movement could be estimated. In order to guarantee that each parameter contributed equally, all data were z transformed prior to analysis.

## Results

### The coupled model

Consumers evolve a larger body size with increasing rates of isolation (Fig 2 & 3). The effect of isolation is additionally strengthened under conditions where resource growth speed is reduced (Fig 2). When isolation is low, adult body size distributions are right skewed with high kurtosis, whereas with increasing isolation, these distributions become more left skewed with low kurtosis (Fig Sl.l & S1.2). Selection of increasing consumer body size with increasing patch isolation is associated with low consumer abundances (Fig S1.3) and rare but far movements (Fig S1.4 & S1.5) relative to metapopulations with highly connected patches. As expected, the level of occupancy decreases with increasing isolation (except for simulations with a growth speed of 0.9 and *NND* of 10; Fig SI.6).

**Figure 2:**
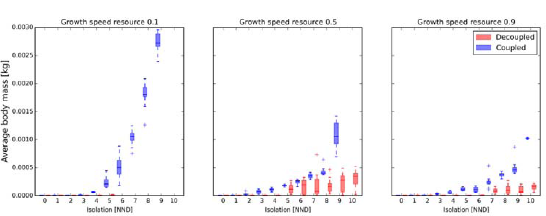
Effect of isolation and resource growth speed on the average adult body mass (*W_max_*) of a consumer. In the coupled model, movement is dependent on body size, while in the decoupled model, both are independent. *NND:* nearest-neighbor distance expressed in number of cells.

**Figure 3:**
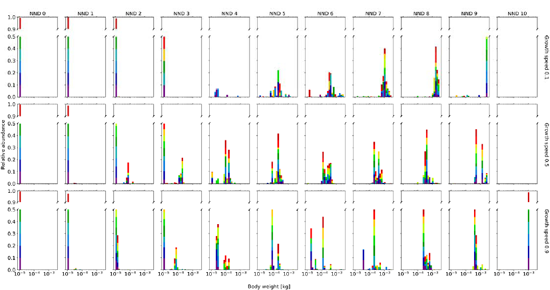
A detailed overview of the evolved adult body size (*W_max_*) distribution of a consumer feeding on a resource when movement is dependent on body size (coupled model). The body size distribution of the consumer clearly depends on the degree of isolation within the landscape (*NND:* Nearest Neighbor Distance) and growth speed of its resource. Per scenario, ten simulations were run. Each simulation is displayed in a different color.

At intermediate levels of isolation and a growth speed of 0.5 or 0.9, body size is often distributed with two peaks within one simulation (see colors in Fig 3). In most simulations, the abundances of these optimal body sizes appear to fluctuate between these two optima, coinciding with a fluctuation in the amount of resources available within the landscape (Fig S1.7). At these intermediate distances, the diversity in size is largest and the coexistence of these multiple body sizes within a community leads to a more efficient depletion of resources within the landscape and lower individual starvation rates (Fig S1.8, S1.9). Population or community-level resource depletion is lowest when nearest neighbor distance is largest (Fig S1.8). When growth speed is low and isolation intermediate (*NND* 3-6), extinction is common during the initial stages of a simulation due to fast depletion of the resources and few reachable patches (Table Sl.l). However, if a population survives this stage and reaches equilibrium, individual starvation chance is lowest (Fig S1.9).

The simulations with a low resource growth speed have the highest *α, β_2_* and *γ* variability (Fig S1.10-12). When growth speed is high, *α* and *β_2_* variability decrease with increasing isolation (Fig S1.10, Sl.ll).

### The decoupled model

Importantly, all simulations with low growth speed (0.1) and high levels of isolation (more than 5 *NND*) go extinct within the decoupled model (Table S1.1). Generally, size-independent movement selects for smaller average adult body sizes (Fig 2). Interestingly, when immigration of novel genotypes within the metapopulation is not allowed (*q=0)*, adult body size converges to the minimum in almost all simulations (p5 sensitivity analysis). In metapopulations with large nearest neighbor distances, this minimum size is not obtained when immigration is allowed (*q=*0.1) (Fig 4). At low and intermediate levels of isolation, the smallest individuals of 0.01 g are being selected (Fig 2 & 4). Therefore, the level of kurtosis and skewness of these simulations are higher than within the coupled model (Fig S1.1, S1.2). Globally, more individuals are present within the decoupled model than the coupled model (Fig S1.3). The number of individuals increases slightly with moderate isolation but decreases drastically at high levels of isolation (Fig S1.3). The average amount of resources shows an opposite pattern (Fig S1.8). Due to the decoupling of body size and movement, individuals move further (Fig S1.4). Simultaneously, the chance of moving during a day is also higher, except when isolation is low (Fig S1.5). As such, the total average distance moved during a lifetime is on average higher within the decoupled model (Fig S1.13). At high levels of isolation, the chance of dying due to starvation is remarkably lower (Fig S1.9), resulting in a longer lifetime (Fig S1.14). Due to changes in movement frequency and distance, the level of occupancy is higher within the decoupled model (Fig S1.6). As within the coupled model, the simulations with a growth speed of 0.1 appear to have the highest α, *β_2_* and *γ* variability (Fig S1.10-12).

**Figure 4:**
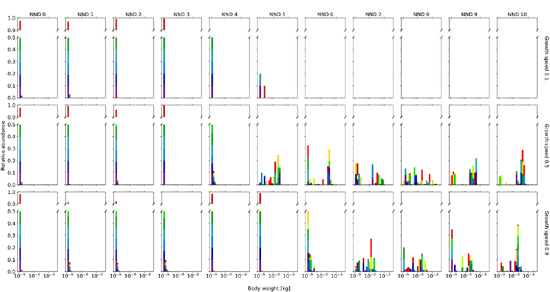
A detailed overview of the evolved optimal adult body size (*W_max_*) distribution of a consumer feeding on a resource when movement is independent of body size (the decoupled model). The effect of isolation (A/A/D: Nearest Neighbor Distance) and growth speed of the resource on the optimal body size distribution of the consumer is shown. Per scenario, ten simulations were run. Each simulation is displayed in a different color.

### Partitioning variance: Importance of size-dependent movement for selection and ecological dynamics

Most variation in adult body size distributions is explained by the level of isolation (Table 1). Growth speed and size-dependent movement are less but almost equally important for the weight distribution of a consumer (Table 1). Growth speed explains most of the total variation in occupancy rate (O), consumer and resource dynamics and metapopulation statistics *a, β_2_* and *γ* variability (Table 1). Moreover, the level of isolation is more important than size-dependent movement for *O* and consumer and resource dynamics. The level of isolation only explains very little variation in *α, β_2_* and *γ* variability (Table 1).

**Table 1.** An overview of the amount of variation in (i) weight distribution (average, coefficient of variation, skewness and kurtosis of *W_max_* distribution), (ii) occupancy, (iii) resource and consumer dynamics (resource abundance, resource variance and consumer abundance) and (iv) metapopulation variability (*α, β_2_* and *γ* variability) that can be explained by the coupling of movement and size, the level of isolation, and resource growth speed.

|  | Coupling | Isolation | Resource growth speed |
| --- | --- | --- | --- |
| Weight distribution | 0.07434 | 0.23488 | 0.09935 |
| Occupancy | 0.10450 | 0.03184 | 0.65737 |
| Resource and<br>consumer dynamics | 0.05374 | 0.10723 | 0.28712 |
| Metapopulation<br>variability | 0.0536 | 0.00069 | 0.36578 |

Of all the statistics of interest, the coupling of size and movement has the largest impact on *O* (Table 1). This coupling is able to explain about 5% of the variation in both resource and consumer dynamics and *a, β_2_* and *y* variability (Table 1).

The amount of variation explained by size-dependent movement is highest at *NND=4*, here reaching 52.65% (Fig 5), and lower at higher and lower levels of patch connectedness (Fig 5). Because metapopulation extinction rate is largest under large nearest neighbor distance for the decoupled model -especially when resource growth is low (Table Sl.l)- and because only surviving metapopulations are integrated in the variance partitioning, less variation in body size is explained by size-dependent movement when isolation is strong (Fig 5). The total amount of variation in body size is additionally higher at high isolation than low or intermediate isolation within the decoupled model (Fig 2 & Fig S1.15).

**Figure 5:**
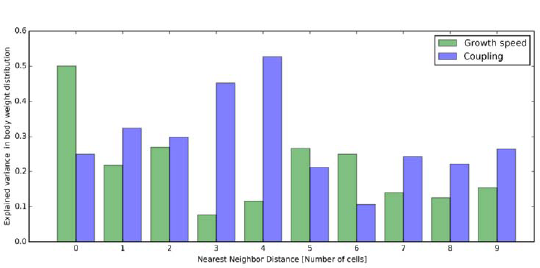
A comparison between the amount of variation in a consumer’s weight distribution that is explained by growth speed and the coupling of body size and movement, for each level of isolation.

The importance of resource growth speed in explaining variation in consumer body size is highest in the most connected landscape (50.09 %) (Fig 4). For the other levels of isolation, size-dependent movement has a higher explaining power than growth speed except for the levels of isolation with *NND* equaling 5 (26.65 % versus 21.20 %) or 6 (24.94 % versus 10.61 %) (Fig 4).

## Discussion

The outcome of our model shows that decreasing connectivity and resource growth select for communities or populations consisting of larger species or individuals due to strong selection for the ability to move over longer distances. Moderate isolation promotes diversity in size, with differently sized individuals able to coexist by foraging at different scales. This increased size diversity also implies higher functional diversity resulting in more efficient resource depletion. Although isolation is the most important driver of consumer body size, resource growth speed is most important for biomass stability, occupancy, and global consumer and resource dynamics. As such, we demonstrate by means of an individual-based model combining metabolic theory and size- dependent or -independent movement that resource productivity and isolation strongly affect the optimal body size distribution. However, especially at intermediate levels of connectivity, size- dependent movement is an important driver of body size distributions and the resulting ecological dynamics and functioning.

### Isolation and resource growth effects on consumer body size distribution and population dynamics

Our results highlight that small-scale isolation has a positive effect on consumer body size. With increasing isolation, larger individuals are selected as only they are capable of crossing unsuitable matrix to reach neighboring patches. This finding is also supported by other theoretical studies (Etienne and Olff 2004). However, experimental research has illustrated that habitat fragmentation can have a variable effect on consumer body size within a population or community (e.g. Sumner, Moritz & Shine 1999; Davies *et al.* 2000; Braschler & Baur 2016). Studies reporting a positive effect contributed this to the positive dependence of mobility on body size (Braschler and Baur 2016, Jauker et al. 2016, Warzecha et al. 2016). With decreasing growth speed of the resource, the positive effect of isolation on consumer body size is amplified. Logically, as fewer resources are available within the landscape, individuals need to move further to locate them, resulting in stronger selection in favor of a larger body size. The effect of resource growth speed and isolation on the weight distribution of the consumer was studied for varying values of the other parameters using a sensitivity analysis (see sensitivity analysis). Although different-sized individuals may move at different relative scales, the general trend of increasing body size with isolation and decreasing resource growth speed is always present. With increasing isolation and decreasing resource growth speed, optimal body size increases, thereby amplifying movement distances, but decreasing movement frequency due to lowered local competition. These effects influence population dynamics by resulting in fewer individuals and lower occupancy levels. As such, consumer populations transform from spatially coupled towards classic metapopulations with increasing isolation and decreasing growth speed of the resource (Amarasekare 2008, Fronhofer et al. 2012).

At intermediate levels of isolation with moderate or high resource growth speed, the body weight distribution of the consumer has two optima. These optima represent a philopatric and a mobile strategy, which coexist and dominate the population depending on the availability of resources. These results suggest that when local resource availability is high, the smaller philopatric individuals flourish, while, once resources are depleted, only the larger, mobile individuals can trace them within the landscape. The fact that these two strategies coexist while feeding on the same resource by foraging at different scales strongly supports the textural discontinuity hypothesis. This hypothesis states that modes of individual size distributions mirror those scales at which resources within the landscape are most abundant (Holling 1992, Laca et al. 2010, Borthagaray et al. 2012, Nash et al. 2014). At intermediate levels of isolation, simulations are characterized by the highest diversity in size. Not coincidently, these simulations are also characterized by being most efficient in resource depletion. Because of niche differentiation along the single axis representing ‘space use’, resources are consumed by each type of body size in a complementing way (Tilman 2001). As such, populations and communities with a higher diversity in size have a higher functional diversity (Song et al. 2014). The observation that size diversity is maximal at intermediate levels of isolation is in line with other studies highlighting a positive effect of fragmentation on species richness (Arnillas et al. 2017, Fahrig 2017).

Within a spatially implicit model and assuming global dispersal, species diversity is optimized at intermediate dispersal rates, increasing spatial insurance of ecosystem functioning (Loreau et al. 2003). We show that when space is considered explicitly, intermediate levels of connectivity select for these intermediate movement distances and rates, which generate high diversity. Although higher functional diversity might imply less variability in overall ecosystem functioning when an abiotic condition is fluctuating in space and time (Isbell et al. 2017), this is not the case in our model. We observe no clear link between size diversity and stability in space (β_2_ variability) or time at the local (*α* variability) or regional (*γ* variability) scale of consumer biomass. Within our model, no extra abiotic condition is included to which consumers can adapt. Still, resources are heterogeneously distributed in space and might fluctuate in time due to consumption and growth. We allow our consumers to adapt to shifts in resource availability by selection of their size and consequently, their behavior and movement. As such, diversity in consumer size results in optimization of resource consumption at intermediate levels of connectivity but not increased stability of consumer biomass.

When isolation is high, resource consumption is decreased. If fragmentation is too strong, this thus leads to non-optimal resource usage at the landscape level, which affects resource availability, an important ecosystem trait.

The lowest number of consumer individuals occurs when resource growth speed is low, resulting in the least stable population dynamics at all three scales (α, *β_2_*, and *γ).* When growth speed is low and isolation intermediate, variability at the regional scale is high, explaining the high number of simulations that went extinct. Surprisingly, α and *β_2_* variability appear to decrease with isolation when growth speed is high. The scenario with both highest isolation and highest growth speed is characterized by high average occupancy but a very low number of individuals, which are large and move rarely but far. This outcome indicates that all these individuals inhabit different cells and are very stationary as resources replenish fast, resulting in the lowest variability of consumer biomass of all scenarios at the local and between-patch scale. This observation indicates the existence of an important interaction between resource productivity and isolation that should be included when studying ecosystem stability. Still, the stability at the regional scale is unaffected by the level of isolation when growth speed is high.

### Importance of size-dependent movement

Two antagonistic forces regulate metapopulation dynamics: selection in favor of short developmental times that increase net growth rate (acting at the within-patch scale) and selection in favor of movement (acting at the between-patch scale) (Davies et al. 2000). Within the coupled model, large individuals are selected when isolation is strong, as only then the benefit of moving far outweighs the disadvantage of developing slowly. However, when decoupling movement speed and body size, an individual’s speed of movement is no longer restricted by its size and instead sampled out of a uniform distribution. As such, the delicate balance between these two forces of selection is disturbed, resulting in generally smaller individuals with fast development rates.

When isolation is high and resource growth speed low, simulations go extinct within the decoupled model as selection for a strategy that guarantees a high movement speed is not possible. Size- dependent movement is thus essential for the survival of actively moving consumer populations and communities when isolation is strong and resource growth speed low. When isolation is high and growth rate moderate or high, resources are more abundant, which enables populations to persist although average movement speeds are lower than within the coupled model. When immigration of novel genotypes into the metapopulation is allowed, these experience a strong advantage when arriving in unoccupied suitable habitat. As such, they increase migration load and strongly influence the population’s average size, which clarifies the large amount of variation in average body size within and between simulations. This migration load also clarifies the high percentage of unexplained variation in weight distribution and corresponding ecological dynamics within the variation partitioning analyses (Table 1). When the immigration of novel genotypes into the metapopulation is deactivated in the decoupled model, smaller body sizes are able to dominate the population or community. Such a decoupling of movement and body size also affects ecological dynamics substantially by resulting in populations and communities with more individuals, which move further and more frequently. Simultaneously, the level of occupancy is increased, which points to spatially coupled populations (Amarasekare 2008, Fronhofer et al. 2012).

Size-dependent movement explains most variation in the consumer’s body size distribution at intermediate levels of connectivity. This contradicts our expectations as we expected size-dependent movement to be most essential for the weight distribution of the consumer at the highest levels of isolation. This is also surprising when considering that the effect of decoupling on average consumer body size is largest at high levels of isolation. However, skewness and kurtosis were least affected by the decoupling at these levels of isolation. Also, the largest individuals are selected in scenarios with low growth speed and high isolation but no comparable simulations could be included of the decoupled model within the analysis as they all went extinct.

The level of isolation has a larger influence on the weight distribution of consumers than the growth speed of the resource or size-dependent movement. Considering that the weight distribution might be interpreted at the community level with each size class representing a different species, these findings support Watling (2006) who states that the importance of isolation for patch richness is expected to increase with ongoing fragmentation of protected areas (Watling and Donnelly 2006).

Body size is central to species vulnerability and functioning. The implementation of our mechanistic model enables a deeper and essential understanding of the impact of fragmentation and altered land use on the organisation of communities and populations. Here, we demonstrate that size-dependent movement is vital for the survival of populations experiencing fragmentation by enabling selection for increased movement speed and therefore larger individuals. Also, size-dependent movement explains most of the observed variation in mass distributions at intermediate levels of connectivity. Further, at these moderate levels of connectivity local size diversity is highest and hence functional diversity, thereby optimising resource control but not stability. Moreover, we highlight an important interaction effect between isolation and resource growth on local and spatial variability. Thereby we contribute to the understanding of the factors that affect body size distributions, enhance size diversity and thereby indirectly affect the stability and functioning of communities across different scales (Isbell et al. 2017).

## Declarations

JH was supported by Research Foundation - Flanders (FWO). The computational resources (STEVIN Supercomputer Infrastructure) and services used in this work were kindly provided by Ghent University, the Flemish Supercomputer Center (VSC), the Hercules Foundation and the Flemish Government - department EWI.

DB, TH and JH conceived the ideas and designed methodology; JH designed the model; DB, MLV, TH and JH analyzed the data; DB, MLV and JH led the writing of the manuscript.

## Data Accessibility

All data will be accessible at github.ugent.be.

